# A microRNA family exerts maternal control on sex determination in *C. elegans*

**DOI:** 10.1101/083949

**Authors:** Katherine McJunkin, Victor Ambros

## Abstract

Gene expression in early animal embryogenesis is in large part controlled post-transcriptionally. Maternally-contributed microRNAs may therefore play important roles in early development. We have elucidated a major biological role of the nematode *mir-35* family of maternally-contributed, essential microRNAs. We show that this microRNA family regulates the sex determination pathway at multiple levels, acting both upstream and downstream of *her-1* to prevent aberrantly activated male developmental programs in hermaphrodite embryos. The predicted target genes that act downstream of the *mir-35* family in this process, *sup-26* and *nhl-2*, both encode RNA binding proteins, thus delineating a previously unknown post-transcriptional regulatory subnetwork within the well-studied sex determination pathway of *C. elegans*. Genome-wide profiling of SUP-26 binding targets reveals 775 mRNAs, most of which have no known role in sex determination, suggesting that the *mir-35* family may modulate numerous other pathways via regulation of *sup-26*. Since sex determination in *C. elegans* requires zygotic gene expression to read the sex chromosome karyotype, early embryos must remain gender-naïve; our findings show that the *mir-35* family microRNAs act in the early embryo to function as a developmental timer that preserves naïveté and prevents premature deleterious developmental decisions.

## Introduction

MicroRNAs are endogenously encoded small RNAs that counteract the translation and stability of complementary mRNA targets (Ketting 2011). Canonical microRNAs (miRNAs) are processed from long primary transcripts by a series of nucleolytic cleavages and ultimately are loaded into Argonaute proteins to form a functional RNA-Induced Silencing Complex, or RISC (Ha and Kim 2014).

Nucleotides 2-8 at the 5’ end of the miRNA are known as the seed sequence because these positions are the most important for target recognition. MiRNAs sharing an identical seed sequence are grouped together as a “family” because they are predicted to redundantly repress a set of common targets. Phenotypic analysis in *C. elegans* and other organisms supports this notion; mutation of all members of a miRNA family generally exhibits a more severe phenotype than deletion of a single miRNA (Alvarez-Saavedra and Horvitz 2010; Ventura et al. 2008; Parchem et al. 2015).

The roles played by miRNAs in embryonic development are incompletely understood. Notable exceptions are the mammalian miR-290 and miR-302 clusters and orthologous miR-430 (zebrafish), which promote pluripotency after the oocyte-to-embryo transition (OET) (Greve et al. 2013; Giraldez 2010). Prior to the OET, gene expression is regulated post-transcriptionally by maternally-contributed factors, and many miRNA families are present during this developmental period (Chiang et al. 2010). However, for most of these maternally-contributed miRNAs, their roles in early development are unknown.

Many prototypical zygotically-expressed miRNAs (e.g. miR-430, *lin-4*, *let-7*) act to control developmental timing, promoting progression to a later developmental stage when their expression is developmentally upregulated. In contrast, maternally-contributed miRNAs are present at the beginning of development and are downregulated as development progresses. Maternally-contributed miRNAs may therefore act in early embryos to enforce a naïve state, preventing developmental decisions from being executed prematurely, for example by dampening gene expression noise, and/or by exerting transient repression of maternal transcripts whose expression is critical later in development. This latter hypothesis is consistent with recent findings that the mechanism of miRNA-mediated target repression in early embryogenesis differs from that observed in later embryos or differentiated tissues by favoring translational repression (which in principle should be reversible), rather than irreversible mRNA decay (Bazzini et al. 2012; Subtelny et al. 2014).

To better understand the biological roles that maternally-contributed miRNAs play in embryonic development, we sought to dissect the genetic network controlled by the nematode-specific *mir-35* family. The *mir-35* family is maternally-contributed as well as zygotically expressed in early embryos, suggesting that it acts at an earlier developmental stage than previously-studied zygotically-expressed animal miRNAs (Alvarez-Saavedra and Horvitz 2010; Wu et al. 2010; Stoeckius et al. 2009). This family is essential for embryonic development in *C. elegans*, but the precise nature of its essential functions are unknown (Alvarez-Saavedra and Horvitz 2010). Besides its essential functions, the *mir-35* family also exhibits pleiotropic effects on diverse embryonic and postembryonic processes, indicating that these miRNAs functionally engage with diverse gene regulatory networks in the embryo (Liu et al. 2011; Massirer et al. 2012; McJunkin and Ambros 2014; Kagias and Pocock 2015).

Here we provide evidence that one of the major roles of the *mir-35* family is to act as a developmental timer, preventing premature and aberrant decision-making in the sex determination pathway. Through a partially maternal effect, the *mir-35* family acts on multiple levels of the sex determination pathway to prevent the spurious activation of male developmental programs. At least two predicted *mir-35* family target genes, *sup-26* and *nhl-2*, both of which encode RNA-binding proteins, act downstream of the *mir-35* family in this pathway. SUP-26 binds a broad range of targets, many of which are unrelated to sex determination, and NHL-2 is known to function in conjunction with diverse miRNAs (Hammell et al. 2009), so these findings widen the scope of pathways potentially regulated by the *mir-35* family through the SUP-26 and NHL-2 axes. We have thus defined a novel regulatory subnetwork consisting of multiple layers of post-transcriptional control that contribute to the timing and fidelity of *C. elegans* sex determination and potentially other embryonic developmental processes.

## Results

### Most transcripts derepressed in *mir-35-41(nDf50)* mutant embryos not direct targets of *mir-35* family miRNAs

The *mir-35* family consists of eight members, *mirs-35-42*, encoded at two loci (Alvarez-Saavedra and Horvitz 2010). The *mir-35-41* cluster encodes seven members of the family on a single transcript, whereas *mir-42* is encoded separately as part of the *mir-42-45* cluster (Figure 1A). Deletion of all eight *mir-35-42* family members results in completely penetrant embryonic lethality, with arrest at various stages of embryogenesis (Alvarez-Saavedra and Horvitz 2010). Forward genetic screens identified no suppressors of *mir-35* family mutant lethality, suggesting that perturbation of multiple unknown pathways contributes to this phenotype (Alvarez-Saavedra and Horvitz 2010).

**Figure 1.**
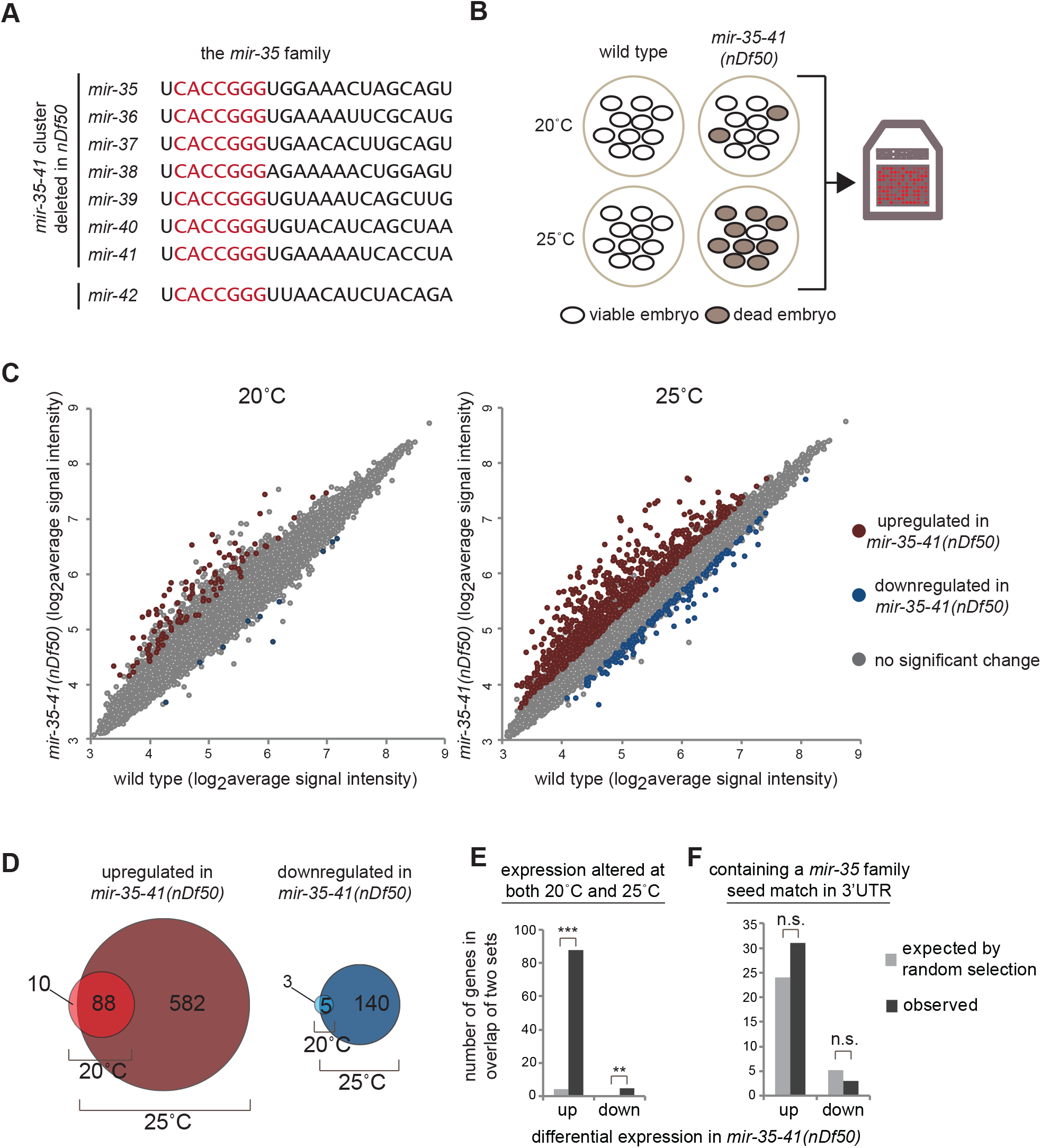
Many genes are upregulated in *mir-35-41(nDf50)* mutants but are not predicted *mir-35* family targets. (A) Sequences of mature microRNAs comprising the *mir-35* family. *mir-35-41* are processed from a single transcript while *mir-42* is located in a separate cluster. Red text indicates the seed sequence. (B) *mir-35-41(nDf50)* is a temperature-sensitive mutant in which embryonic lethality is low at 20°C and highly penetrant at 25°C. Gene expression in embryos raised at each temperature was profiled using microarrays. (C) Many more genes are differentially expressed in *mir-35-41(nDf50)* embryos at 25°C than at 20°C. More genes are upregulated than are downregulated. (D) The genes that are differentially expressed at 20°C are largely a subset of those observed at 25°C. (E) The sets of differentially expressed genes at 20°C and at 25°C overlap much more than expected by chance. ***, p-value < 1x10^-100^. **, p-value < 1x10^-5^ (hypergeometric test). (F) The genes differentially expressed in *mir-35-41(nDf50)* embryos do not contain more *mir-35* family seed matches than expected in a randomly chosen gene set.

Deletion of the *mir-35-41* cluster alone (leaving *mir-42* intact) yields a hypomorphic and temperature-sensitive phenotype (Alvarez-Saavedra and Horvitz 2010). *Mir-35-41(nDf50)* mutants exhibit incompletely penetrant embryonic lethality, the penetrance of which increases with temperature. This malleable genetic setting has enabled the characterization of numerous post-embryonic and pleiotropic phenotypes of *mir-35* family loss of function (Massirer et al. 2012; Liu et al. 2011; McJunkin and Ambros 2014; Kagias and Pocock 2015). Still, a clear molecular basis for the phenotypes of *mir-35* family mutants has remained elusive. We took advantage of this temperature-sensitive allele (*mir-35-41(nDf50)*) to characterize the molecular phenotype associated with *mir-35* family loss of function and to identify downstream target genes responsible for the phenotype.

To gain insight into the molecular phenotype caused by *mir-35* family loss of function, we performed gene expression analysis of *mir-35-41(nDf50)* mutant embryos. Since the *nDf50* deletion allele acts as a temperature-sensitive, hypomorphic allele of *mir-35* family function, we profiled gene expression changes at both the semi-permissive (20°C) and restrictive (25°C) temperatures (Figure 1B).

Consistent with the increased severity of phenotypes at restrictive temperature, many more gene expression changes were observed in embryos raised at 25°C (670 genes) than in those raised at 20°C (98 genes) (cutoff >1.5-fold, p-value < 0.05, Figure 1C, Table S1). The small set of deregulated genes observed at 20°C was nearly an exact subset of the differentially expressed genes at 25°C (Figure 1D-E). This supports the notion that the phenotypes observed at restrictive temperature are a more severe manifestation of the mild phenotypes observed at permissive temperature.

At both 20°C and 25°C, the genes upregulated in *mir-35-41(nDf50)* mutants compared to wild type greatly outnumbered those downregulated in the mutants (98 versus 8 at 20°C and 670 versus 145 at 25°C) (Figure 1C-D). We postulated that the large number of upregulated genes might represent direct targets of the *mir-35* family miRNAs, which are derepressed in the miRNA mutants. This was not the case. The set of upregulated genes was not enriched for genes containing *mir-35* family seed matches in their 3’UTRs (Figure 1F). Instead, the number of genes predicted to be targeted by the *mir-35* family in this set was similar to the number expected by chance. The genes downregulated in *mir-35-41(nDf50)* mutant embryos were likewise not enriched for *mir-35* family seed matches. Therefore, the majority of the gene expression changes observed in *mir-35-41(nDf50)* mutant embryos represent downstream, indirect effects of *mir-35* family loss of function.

### *mir-35-41* prevents masculine development at multiple levels in the sex determination pathway

Previously, we demonstrated that a predicted *mir-35* family target gene implicated in the sex determination pathway, *SUPpressor-26 (sup-26),* is epistatic to fecundity phenotypes observed in *mir-35-41(nDf50)*. We wondered whether *mir-35-41(nDf50)* mutants might have a previously undetected sex determination phenotype. Therefore, we asked whether the molecular phenotype, i.e. gene expression changes, of *mir-35-41(nDf50)* mimicked those of a sex determination mutants by examining published datasets.

We observed a striking correlation between the gene expression changes in *mir-35-41(nDf50)* mutants and those in loss-of-function mutants of *sdc-2* (sex determination and dosage compensation defect 2) (r=0.49, p-value<.0001) (Figure 2A) (Jans et al. 2009). In particular, the derepressed genes in each genotype were highly overlapping and strongly correlated (Figure 2A-B).

**Figure 2.**
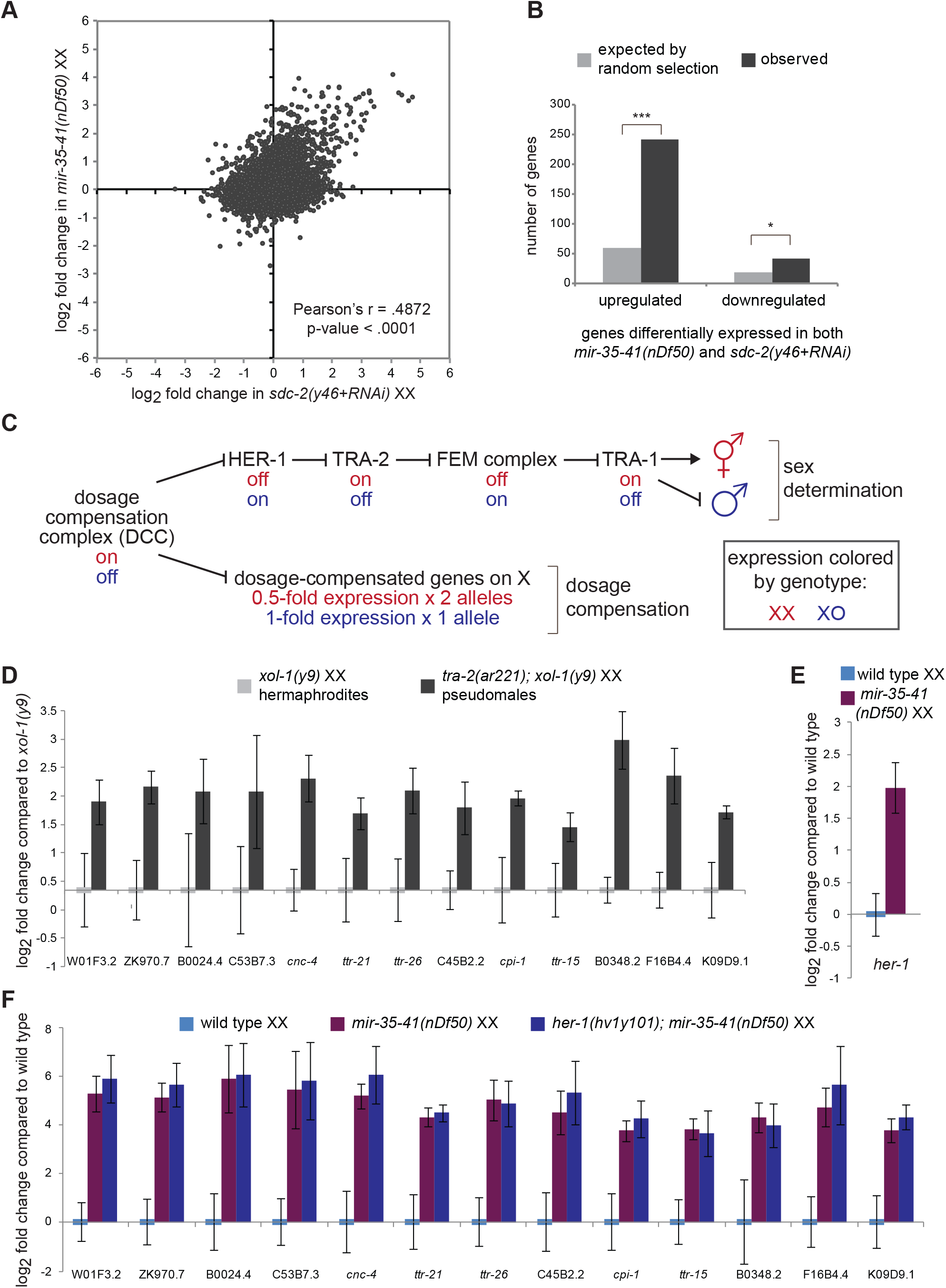
Gene expression changes in *mir-35-41(nDf50)* mutants are similar to those in sex determination mutants. (A) Comparison of gene expression changes in *mir-35-41(nDf50)* embryos and *sdc-2(lf)* embryos, each normalized to wild type. (B) The set of genes differentially regulated in *mir-35-41(nDf50)* and in *sdc-2(lf)* overlaps more than expected by chance. ***, p-value < 1 x 10^-80^. *, p-value < 1 x 10^-5^ (hypergeometric test). (C) Genetic model of *C. elegans* sex determination and dosage compensation. (D-F) qRT-PCR of transcripts indicated on X-axis in embryos of genotypes indicated above graph. The mean and SEM of three biological replicates is shown. (D) Both genotypes are normalized to *xol-1(y9)*. (E-F) All genotypes are normalized to wild type XX embryos.

*Sdc-2* encodes a core component of the dosage compensation complex (DCC), a condensin-like transcriptional repressor that carries out two functions. One function of the DCC is to bind along the X chromosome to downregulate gene expression by two-fold (dosage compensation). Since the DCC is active only in XX animals after the 40-cell embryo stage due to karyotype-specific expression of *sdc-2*, this function equalizes X-linked gene dosage between XX hermaphrodites and XO males. The second function of the DCC is to control sex determination by repressing the master driver of male sexual fate, *her-1* (hermaphroditization-1, an autosomal gene) by 40-fold, leading to hermaphroditic sexual development in XX animals (Meyer 2005) (Figure 2C). Because of the DCC’s role in both processes, XX animals in which DCC function is impaired display lethality due to loss of dosage compensation (DC) but also aberrantly masculinized development (Tra, Transformer phenotype) due to derepression of *her-1*.

Many of the upregulated genes common to *mir-35-41(nDf50)* and *sdc-2(lf)* were not X-linked and were upregulated greater than two-fold, suggesting that these observed gene expression changes are reflective of perturbed sex determination (a molecular Tra phenotype) rather than derepressed targets of DC. In further support of this notion, *mir-35-41(nDf50)* mutant gene expression changes did not significantly overlap with those in a mutant that disrupts only DC but not sex determination, *dpy-27(y57)* (Jans et al. 2009).

To further test whether the gene expression changes in *mir-35-41(nDf50)* were due to an aberrantly activated male gene expression program, we generated masculinized (Tra) embryos using an orthogonal genetic means: a mutation in the sex determination pathway (*tra-2(ar221);xol-1(y9)*). If the high-amplitude gene expression changes observed in *mir-35-41(nDf50)* and *sdc-2(lf)* are symptomatic of masculinization, then similar changes in gene expression should be observed in the pseudomale embryos generated by perturbing the sex determination pathway directly. This was indeed the result: the thirteen genes that were most highly upregulated in *mir-35-41(nDf50)* embryos are similarly upregulated in *tra-2(ar221);xol-1(y9)* pseudomale embryos (Figure 2D). Therefore, many of the measurable gene expression changes in *mir-35-41(nDf50)* embryos are symptomatic of aberrantly activated masculine development. We term this phenotype pseudomale gene expression (ψGE), and use this panel of thirteen genes as a readout of the ψGE phenotype.

To determine the level at which the sex determination pathway is perturbed in *mir-35-41(nDf50)* mutants, we examined *her-1* transcript levels by Q-PCR. Elevation of *her-1* was observed in *mir-35-41(nDf50)* embryos, indicating that the sex determination pathway is deregulated upstream of *her-1* (Figure 2E). Surprisingly, the broader ψGE program in *mir-35-41(nDf50)* mutant XX embryos is not a direct consequence of upregulated *her-1*: ψGE was still observed in *mir-35-41(nDf50)*; *her-1(hv1y101null)* double mutants (Figure 2F). This indicates that signaling in the sex determination pathway is aberrantly masculinized both upstream and downstream of *her-1* in *mir-35-41(nDf50)*. Therefore, the *mir-35-41* family acts both upstream of *her-1* to indirectly repress *her-1* transcription and downstream of *her-1* to prevent the activation of a broader male gene expression program (ψGE).

### *mir-35-41(nDf50)* mutants are cryptically masculinized

Despite the activation of *her-1* transcription and the ψGE program, sex determination of *mir-35-41(nDf50)* XX hermaphrodites appears superficially normal. We hypothesized that *mir-35-41(nDf50)* XX embryos might be cryptically masculinized, and that this might be revealed by the appearance of overt masculinization in a sensitized genetic background. Accordingly, we utilized the *her-1(n695gf)* allele, which causes a weak derepression of *her-1* transcription in XX animals, leading to mild masculinization in hermaphrodites, evidenced by an egg-laying defective (Egl) phenotype. We crossed *mir-35-41(nDf50)* into *her-1(gf),* and observed that *mir-35-41(nDf50);her-1(gf)* double mutants exhibit dramatically enhanced synthetic masculinization (synTra phenotype); nearly all doubly-mutant XX animals are self-sterile pseudomales (Figure 3A-B). This synTra phenotype is rescued by expression of *mir-35-45* from an extrachromosomal array (Figure 3B). Thus, deletion of *mir-35-41* strongly enhances masculinization of *her-1(gf)*, and *mir-35-41* therefore promotes hermaphroditic sex determination in wild type XX animals.

**Figure 3.**
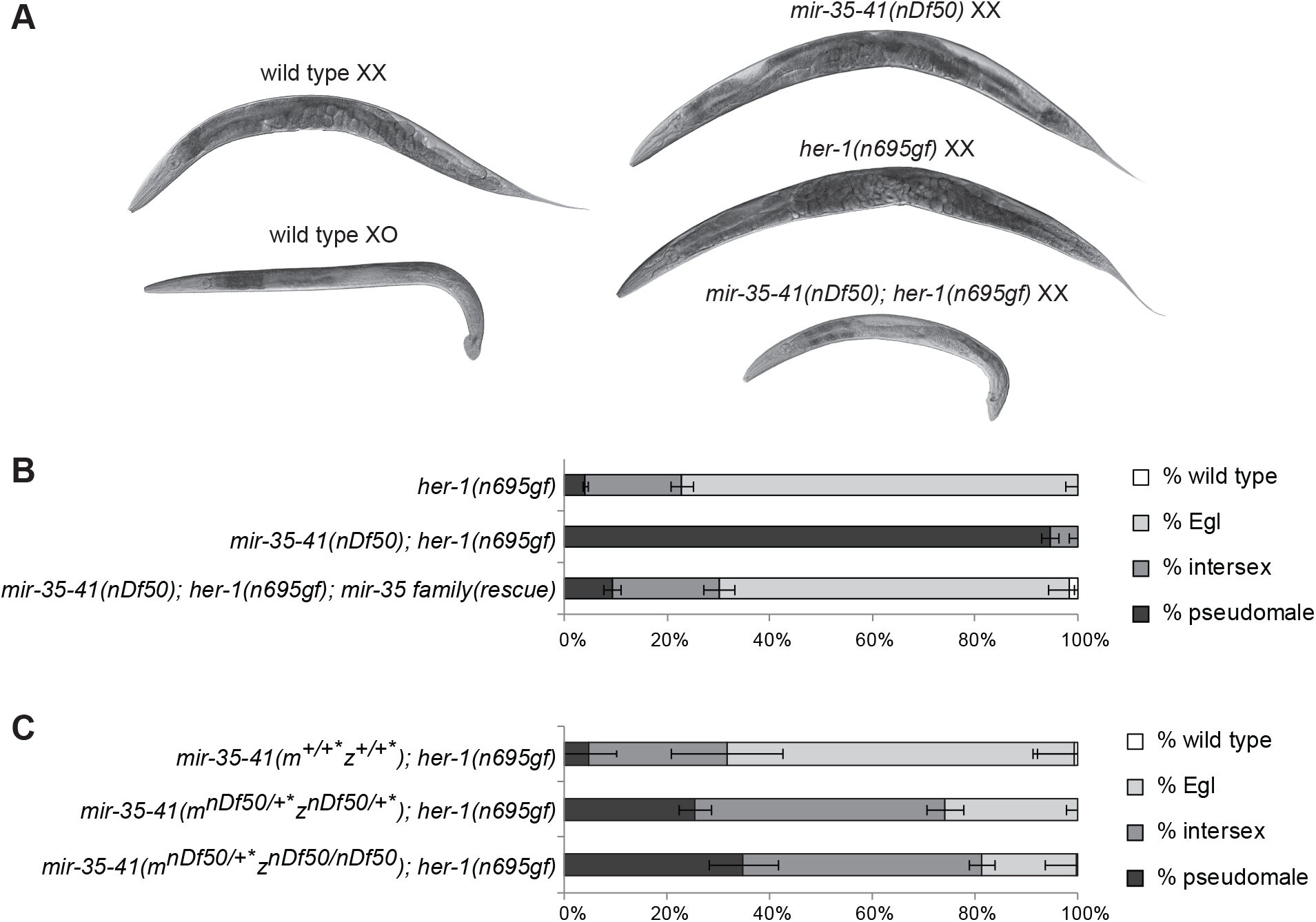
*mir-35-41* are required for proper sex determination in hermaphrodites. (A) DIC micrographs of wild type and *her-1(gf)* and *mir-35-41(nDf50)* single and double mutants. (B) Quantification of sex determination phenotypes; darker colored bars indicate more severe masculinization. (C) The requirement for *mir-35-41* in sex determination is dose-dependent since heterozygotes display enhanced masculinization. Removing both copies of zygotic *mir-35-41* does not strongly further enhance masculinization, indicating a strong requirement for the maternal contribution of *mir-35-41*. +*, mIn1 balancer chromosome.

Interestingly, sex determination is highly sensitive to the dosage of *mir-35-41*. Heterozygotes for *mir-35-41(nDf50)* also show significant enhancement of *her-1(gf)* masculinization (Figure 3C). Furthermore, the maternal contribution of *mir-35-41* is crucial for sex determination; animals whose mothers contain one copy of the cluster show a similar sex phenotype regardless of whether they inherit a zygotic copy of *mir-35-41* (Figure 3C, bottom two bars). Moreover, in animals lacking zygotic *mir-35-41*, a single maternal copy of *mir-35-41* rescues pseudomale penetrance from 94% to 35% (compare middle bar of Figure 3B to bottom bar of Figure 3C). This strong maternal effect suggests that the *mir-35* family can function at the earliest stages of embryonic development to prevent male development of XX animals.

### The predicted *mir-35* family target genes *sup-26* and *nhl-2* are required for cryptic masculinization of *mir-35-41(nDf50)*

Neither *her-1* nor other genes in the core sex determination pathway are predicted targets of the *mir-*35 family, based on searches for *mir-35* seed matches in 3’ UTRs. Therefore, we sought to determine which direct *mir-35* family target genes are responsible for the masculinized gene expression signature and cryptic masculinization in *mir-35-41(nDf50).* To this end, we screened 72 predicted *mir-35* family target genes by RNAi to assess which genes were required for the synTra phenotype of *mir-35-41(nDf50);her-1(gf)* animals (Table S2). Of these genes, two were each required for the synTra phenotype: *sup-26* and *nhl-2* (*NHL* domain-containing). The effects of RNAi were recapitulated using mutant alleles of each gene. While *mir-35-41(nDf50);her-1(gf)* animals develop as pseudomales, *mir-35-41(nDf50);her-1(gf);sup-26(lf)* or *mir-35-41(nDf50);her-1(gf);nhl-2(lf)* develop as wild type or Egl hermaphrodites (Figure 4A-B). Thus, the predicted target genes *sup-26* and *nhl-2* are epistatic to the enhanced masculinization caused by *mir-35-41(nDf50)* and likely act downstream of *mir-35-41*.

**Figure 4.**
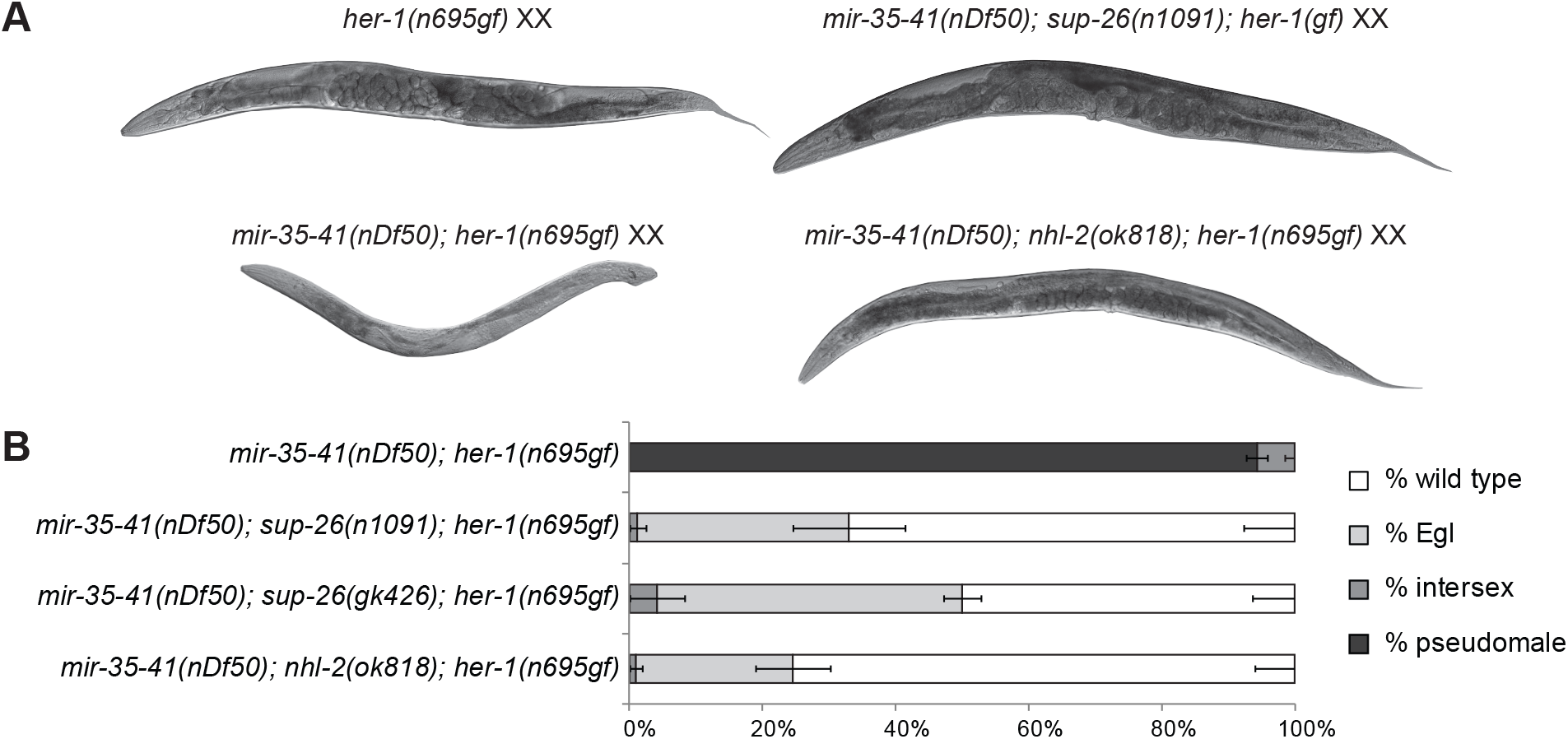
The *mir-35* family target genes *sup-26* and *nhl-2* are required for the sex determination phenotypes of *mir-35-41(nDf50).* (A) Representative DIC micrographs of indicated genotypes. (B) Quantification of sex determination phenotypes; darker colored bars indicate more severe masculinization.

*sup-26* encodes an RNA-binding protein (RBP) containing a Q-rich low complexity domain and two RNA recognition motifs (Figure 5A). *sup-26* was previously isolated in a screen for suppressors of the Egl phenotype of *her-1(gf)*, and we confirmed this genetic interaction (Manser et al. 2002) (Figure S1). We also previously demonstrated that *sup-26* is required for another aspect of the *mir-35-41(nDf50)* mutant phenotype: temperature-sensitive loss of fecundity (McJunkin and Ambros 2014). To supplement existing alleles of *sup-26,* we generated a null allele using CRISPR/Cas9 genome editing to replace the entire SUP-26 coding sequence with the GFP coding sequence (Figure S2A). This allele behaved similarly to other *sup-26* loss-of-function alleles with respect to *her-1(gf)* and *mir-35-41(nDf50);her-1(gf)* phenotypes (Figure S2B), though the latter could only be tested in *mir-35-41(nDf50)* heterozygotes due to a synthetic lethality of *sup-26(lf) mir-35-41(nDf50)* homozygotes. (The synthetic lethality of *mir-35-41(nDf50)*; *sup-26(lf)* double mutants is counterintuitive, given that *sup-*26*(lf)* suppresses other *mir-35-41(nDf50)* phenotypes; although the nature of this synthetic lethal phenotype is poorly understood, it is nevertheless apparent that the lethality is likely unrelated to the sex determination function of *sup-26* downstream of *mir-35* family.) No gene expression changes were observed in *sup-26(null)* XX embryos, but *sup-26(null)* males exhibited a mating defect, consistent with the role of *sup-26* in promoting male sex determination (Figure S2C-D).

**Figure 5.**
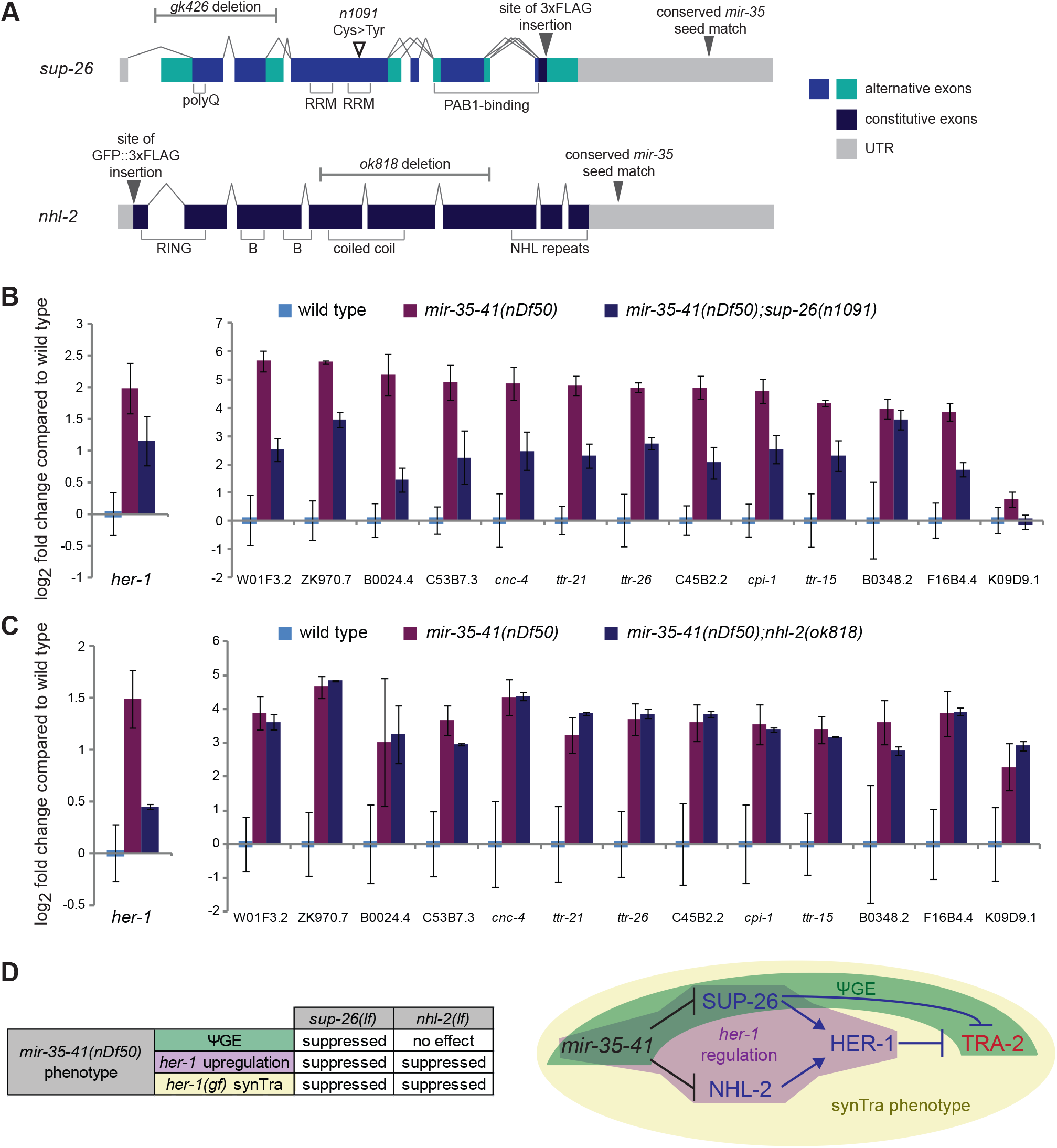
SUP-26 and NHL-2 act at distinct levels in the sex determination pathway. (A) Gene models of *sup-26* and *nhl-2* depicting exon structure, protein domains, mutant alleles and CRISPR-Cas9 editing sites. (B-C) qRT-PCR in embryos raised at 20°C. The mean and SEM of three biological replicates is shown. All genotypes are normalized to wild type XX embryos. (D) Left: Summary of genetic interactions of *mir-35-41(nDf50)* with *sup-26(lf)* or *nhl-2(lf)*. Right: Genetic model highlighting pathways whose disruption in *mir-35-41(nDf50)* mutants underlies each aspect of the mutant sex determination phenotype.

Previous studies have supported a role for SUP-26 in repressing translation of *transformer-2 (tra-2)*, which encodes the target receptor for the HER-1 ligand (Mapes et al. 2010) (Figure 2C). If the function of SUP-26 downstream of *mir-35-41* is entirely at the level of *tra-2*, then the *mir-35-41(nDf50)* ψGE phenotype should be normalized in *mir-35-41(nDf50);sup-26(lf)* embryos, but *her-1* transcription (which is largely controlled upstream of *tra-2*) should remain elevated. Instead *sup-26(lf)* alleviates both the upregulation of the ψGE and that of *her-1* (Figure 5B). Therefore, *sup-26* may act upon the sex determination pathway at multiple levels downstream of *mir-35-41* or via a feedback loop through which sex determination can impact the DCC and hence *her-1* expression (Hargitai et al. 2009).

*nhl-2* encodes a protein of the TRIM (tripartite-containing motif; RING, B-box, coiled-coil domain) −NHL(NCL-1 HT2A and LIN-41 repeat) family, homologous to Mei-P26 and Brat in *Drosophila melanogaster* (Hyenne et al. 2008; Hammell et al. 2009) (Figure 5A). NHL-2 plays a role in polarity of the early *C. elegans* embryo, similar to Brat, which also contributes to cell polarity (Hyenne et al. 2008). NHL-2 promotes the activity of miRNAs in post-embryonic development, while *Drosophila* Mei-P26 represses miRNA abundance (Neumüller et al. 2008; Hammell et al. 2009). Our data, particularly the epistasis of *nhl-2* to *mir-35-41* and the presence of a conserved *mir-35* family seed match in the *nhl-2* 3’UTR, support a novel role for *nhl-2* downstream of *mir-35-41* as a target mRNA.

NHL-2 has not previously been implicated in the DC or sex determination pathways. Like *sup-26(lf)*, *nhl-2(lf)* also suppresses the Egl phenotype of *her-1(gf)* and the derepression of the *her-1* transcript in *mir-35-41(nDf50)* embryos (Figure 5C and Figure S1). The suppression of *her-1(gf)* is particularly surprising since *nhl-2* was not isolated in mutagenesis screens for such suppressors. Unlike *sup-26(lf)*, *nhl-2(lf)* did not suppress the *mir-35-41(nDf50)* ψGE phenotype (Figure 5C). Therefore, *nhl-2* acts upstream of *her-1* in modulating sex-specific development downstream of *mir-35-41*.

Interestingly, these data indicate that the synTra phenotype of *mir-35-41(nDf50);her-1(gf)* requires the elevation of *her-1* itself, and not the ψGE program alone. Since none of the mutants examined here disrupt only the ψGE but not *her-1* upregulation, we cannot conclude whether the ψGE is also required for the synTra phenotype, and we postulate that is does contribute alongside *her-1* upregulation (Figure 5D).

While *sup-26* and *nhl-2* are both required for the sex determination phenotypes of *mir-35-41(nDf50)* mutants, neither gene is required for the embryonic lethality phenotype of these mutants. On the contrary, deletion of *sup-26, nhl-2*, or both genes paradoxically enhances the lethality of *mir-35-41(nDf50)* (Figure S3). This is consistent with the notion that perturbation of multiple pathways likely contributes to *mir-35* family mutant lethality and does not support nor refute a role for *sup-26* or *nhl-2* in contributing to this lethality.

### SUP-26 and NHL-2 localize to P granules in early embryos

Our model is that translational derepression or stabilization of both *sup-26* and *nhl-*2 in *mir-35-41(nDf50)* mutant embryos together contribute to the sex determination phenotypes observed (Figure 5D). Both *sup-26* and *nhl-2* have strikingly long 3’UTRs, 1163-nt and 1151-nt, respectively, compared to the median 3’UTR length of 130-140nt in *C. elegans* (Jan et al. 2011; Mangone et al. 2010), suggesting that these transcripts are subject to post-transcriptional regulation. Both *nhl-2* and *sup-26* (previously known as *tag-310*) are within the top six predicted targets of the *mir-35* family on Targetscan due to a well-conserved perfect 8-mer seed match to the *mir-35* family located in each of their 3’UTRs (Jan et al. 2011). Both mRNAs are also high stringency predicted targets on mirWIP and were identified in sequencing of RNAs physically associated with RISC (Hammell et al. 2008).

Because *mir-35-41* exerts a partially maternal effect on sex determination, we hypothesized that *sup-26* and *nhl-2* might be expressed and regulated by *mir-35-41* in the very early embryo. To examine SUP-26 and NHL-2 abundance in early embryos, we used CRISPR/Cas-9-mediated genome editing to tag SUP-26 and NHL-2 at their endogenous genomic loci (Figure 5A). Three FLAG tags were fused to the C terminus of SUP-26 (SUP-26::FLAG), and GFP and three FLAG tags were fused to the N terminus of NHL-2 (GFP::NHL-2).

Consistent with a role in the early embryo and previous reports of NHL-2 expression, both SUP-26::FLAG and GFP::NHL-2 were expressed in the maternal germline and maternally contributed to early embryos (Figure 6A-B, Figure S4A-B) (Hyenne et al. 2008). Both tagged proteins preferentially localize to perinuclear bodies by the 24-cell stage; these were confirmed to be germline P granules by co-staining for SUP-26::FLAG or endogenous NHL-2 with P granule markers (Figure 6C-D). P granule enrichment is post-transcriptionally controlled since reporters expressing GFP under the control of *sup-26* or *nhl-2* regulatory elements do not show such a bias (Figure S4C) (McJunkin and Ambros 2014). Both SUP-26::FLAG and GFP::NHL-2 are ubiquitously expressed and cytoplasmically localized in later stages of embryonic development, though GFP::NHL-2 remains P lineage-enriched (Figure 5A-B).

**Figure 6.**
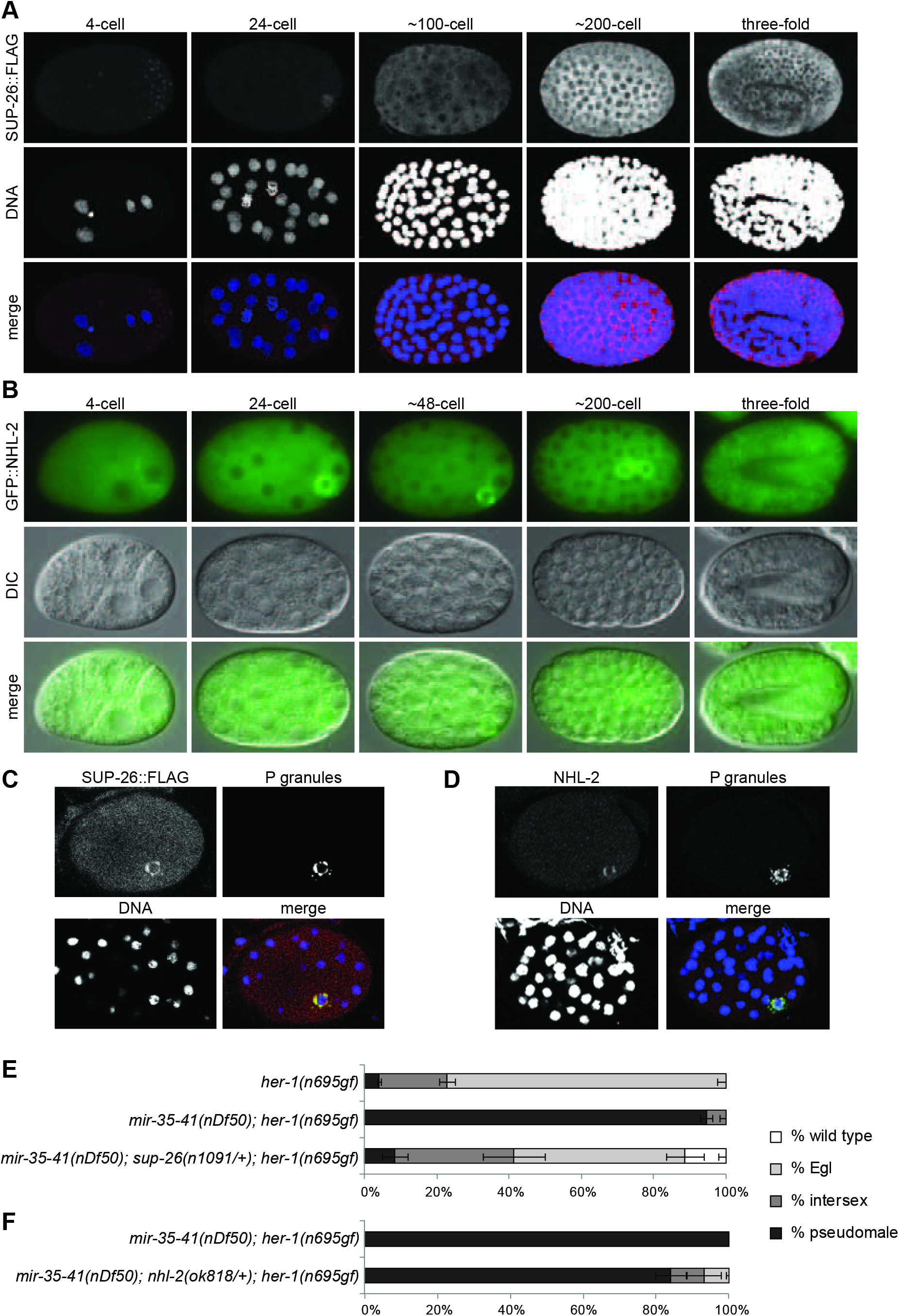
SUP-26 and NHL-2 localize to P granules in early embryos. (A) Maximum projection of confocal stacks showing immunofluorescence of SUP-26::FLAG. (B) Epifluorescence and corresponding DIC images of embryos expressing GFP::NHL-2. (C) Single confocal image of SUP-26::FLAG co-staining with PGL-1, a core component of P granules, in early embryos. (D) Maximum projection of confocal stack of NHL-2 co-staining with a P granule-specific antibody (K76) in early embryos. (E-F) The roles of *sup-26* and *nhl-2* in sex determination are dose-dependent: (E) *sup-26(n1091)* heterozygosity strongly suppress the *mir-35-41(nDf50);her-1(gf)* synTra phenotype. + denotes qC1 balancer chromosome. (F) *nhl-2(ok818)* heterozygosity weakly suppress the synTra phenotype. All animals of both genotypes contain one copy of the qC1 balancer chromosome.

Our model predicts that SUP-26::FLAG and GFP::NHL-2 abundance would be increased in the absence of *mir-35-41.* Surprisingly, we did not observe a robust upregulation of SUP-26::FLAG and GFP::NHL-2 signals in the *mir-35-41(nDf50)* mutant background, though we observed a weak, statistically insignificant trend toward upregulation in early embryonic P granule-localized SUP-26::FLAG. This is especially surprising due to the high seed pairing stability of *mir-35* family members and previous reports of seed match-dependent silencing of a heterologous reporter by the *sup-26* 3’UTR (Garcia et al. 2011; Kagias and Pocock 2015).

We reasoned that *sup-26* and *nhl-2* might be regulated in a highly dose-dependent manner in the endogenous setting, in light of the fact that *mir-35-41* has a strongly dose-dependent effect on sex determination phenotypes (Figure 3C). If this is the case, then SUP-26 and NHL-2 may contribute to the *mir-35-41(nDf50)* sex determination phenotypes through relatively small changes in abundance. Supporting this notion, the effect of *sup-26* on the synTra phenotype of *mir-35-41(nDf50); her-1(gf)* is strongly dose-dependent; heterozygosity of *sup-26(n1091lf)* is sufficient to suppress the phenotype (Figure 6E). *nhl-2* heterozygosity also shows dose-dependent suppression of the *mir-35-41(nDf50);her-1(gf)* phenotype though to a lesser extent (Figure 6F). Thus, the magnitude of SUP-26 and NHL-2 derepression necessary for the sex determination phenotypes of *mir-35-41(nDf50)* is on the order of two-fold, which may be below the sensitivity of our detection method. An alternative model consistent with the data is that SUP-26 and NHL-2 act downstream of *mir-35-41* indirectly, rather than as direct target genes.

### SUP-26 is a promiscuous 3’UTR binding protein whose targets include *nhl-2*

We set out to define the complement of SUP-26 binding targets for two reasons. (1) Previous genetic and biochemical data support a role for SUP-26 in repressing *tra-2* translation to regulate sex determination (Manser et al. 2002; Mapes et al. 2010). However, we observed that *sup-26* was required for the change in *her-1* abundance in *mir-35-41* mutant embryos, suggesting SUP-26 may regulate additional targets upstream of *tra-2.* (2) Any additional targets of SUP-26 binding might also be indirectly regulated by the *mir-35* family and contribute to other aspects of the *mir-35* family mutant phenotype.

To this end, we confirmed that the FLAG tags did not disrupt the wild type function of SUP-26, and performed high throughput sequencing coupled with crosslinking immunoprecipitation (HITS-CLIP) using the SUP-26::FLAG strain (Figure S5A-B). Radiolabeling of RNA in CLIP samples showed a strong signal in crosslinked SUP-26::FLAG samples, but no signal in samples lacking SUP-26::FLAG or in non-crosslinked SUP-26::FLAG samples (Figure S5C). This indicates that the cloned RNA was directly crosslinked to SUP-26::FLAG.

SUP-26::FLAG HITS-CLIP yielded 857 peaks which were detected in at least three out of four biological replicates (Table S3). Analysis of the peak locations revealed that >95% of peaks overlapped with the 3’UTR of 775 protein-coding genes (Figure 7A). Thus, SUP-26 is clearly a 3’UTR binding protein with a large repertoire of targets. Functional annotation of transcripts bound by SUP-26 revealed that the targets are enriched in genes involved in embryonic and larval development, consistent with the strong expression of SUP-26 during these stages (Figure 7B). The proportion of SUP-26::FLAG target transcripts that contain a *mir-35* family seed match is only slightly higher than in the genome at large (5.9% versus 3.2%), confirming the notion that SUP-26 and *mir-35-42* for the most part do not act in concert.

**Figure 7.**
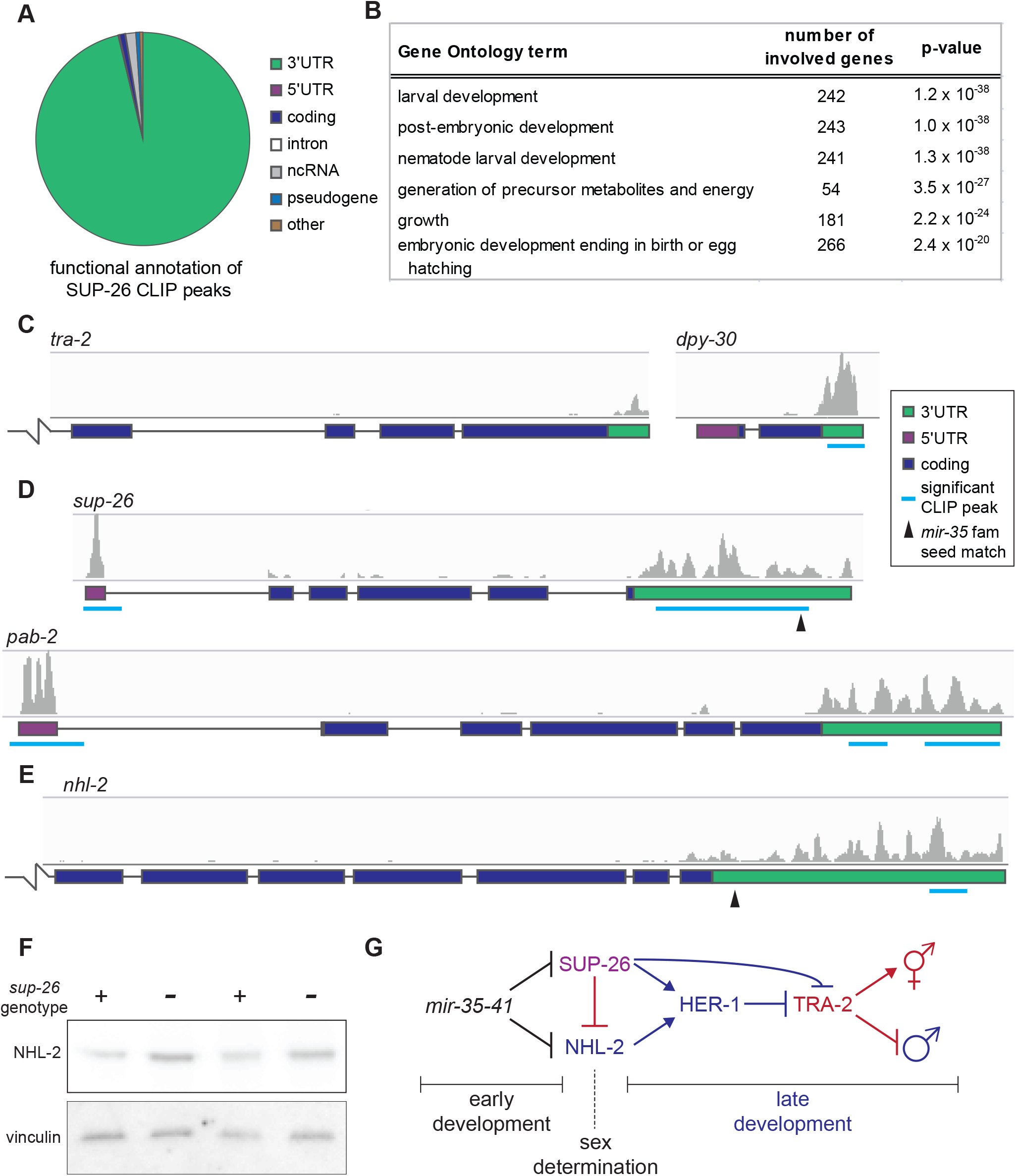
SUP-26 binds numerous 3’UTRs, including that of *nhl-2*. (A) Functional annotation of significant SUP-26::FLAG CLIP peaks. (B) Top significantly enriched gene ontology terms among transcripts bound by SUP-26::FLAG. (C-E) Example CLIP traces showing SUP-26::FLAG binding to (C) the 3’UTR of transcripts implicated in sex determination and dosage compensation (*tra-2* and *dpy-30*), (D) the 5’UTR and 3’UTR of *sup-26* and *pab-2*, and (E) the 3’UTR of *nhl-2*. Broken line indicates that gene model continues outside the frame. (F) Western blot showing upregulation of NHL-2 in *sup-26(null)* mutants. Two biological replicates are shown. (G) Simplified working model of the *mir-35-41* regulatory network. *mir-35-41* likely acts in early development in both XX and XO embryos to repress *sup-26* and *nhl-2* prior to sex determination. *sup-26* and *nhl-2* promote male development, and may indirectly regulate HER-1 by affecting DCC activity (not shown).

CLIP signal corresponding to SUP-26 binding was observed in the 3’UTR of SUP-26’s previously-described target, *tra-2*, though the signal was not identified as a significant peak (Figure 7C). Another gene that could also contribute to the role of SUP-26 in sex determination, *dpy-30*, displayed stronger SUP-26 binding (Figure 7C); *dpy-30* encodes a core member of the DCC. Many other genes that are not implicated in sex determination or DC also displayed strong SUP-26 binding in their 3’UTRs (Figure S5D). A significant sequence motif was not identified in SUP-26 CLIP peaks, suggesting that the specificity of SUP-26 binding may be largely determined by factors other than primary sequence. Overall, these data indicate that SUP-26, and in turn the *mir-35* family, may have widespread effects on translation during development.

Interestingly, SUP-26 also binds the 5’UTR of two genes: its own transcript and that encoding poly(A)-binding protein 2 (*pab-2)* (Figure 7D). This broad repertoire of binding to 3’UTRs but highly specific 5’UTR binding is reminiscent of cytoplasmic poly(A)-binding proteins (*pab-1* and *pab-2* in *C. elegans*), which regulate the translation of their own transcript via 5’UTR binding (Mangus et al. 2003). While SUP-26 does not appear to bind poly(A) *per se*, it was previously shown to exist in a complex with PAB-1 (Mapes et al. 2010) (Figure 5A). We propose that SUP-26 may act with PAB-1 and PAB-2 to regulate translation and feed back on transcripts encoding members of this complex to tightly regulate the complex’s abundance.

The newly-defined targets of SUP-26 binding include *nhl-2*. A significant CLIP peak was called in the distal region of the *nhl-2* 3’UTR, nonoverlapping with the *mir-35* family seed match (Figure 7E). Western blots in mixed-stage embryo samples show that NHL-2 is upregulated in *sup-26(null)* embryos, suggesting that SUP-26 binding of *nhl-2* may repress its translation (Figure 7F). This interaction may contribute to the failure to detect changes in abundance in GFP::NHL-2 in *mir-35-41(nDf50)*; the derepression of *gfp::nhl-2* from *mir-35* family repression may be dampened by simultaneous derepression of *sup-26* (Figure 7G). In general, the SUP-26 HITS-CLIP data suggests that the regulatory network controlling sex determination downstream of the *mir-35* family is highly interconnected and may feed back on itself at multiple levels.

## Discussion

Here we show that a major function of the maternally-contributed *mir-35* family is to contribute to the timing and fidelity of sex determination in *C. elegans*. This is the first demonstration of a role for miRNAs in the *C. elegans* sex determination pathway. Our data show that the predicted *mir-35* family target genes *sup-26* and *nhl-2* act downstream of the *mir-35* family in controlling this process. Taken together, our study of this miRNA family and its targets has delineated multiple new layers of post-transcriptional control (via both miRNAs and RBPs) over the sex determination pathway.

In contrast to most players in the sex determination pathway, the *mir-35* family has a strong maternal effect. Most genes in the sex determination act zygotically, only after the embryo’s X to autosome ratio is assessed, via a process of monitoring zygotic transcription of competing X-linked and autosomal signal elements (around the 40-cell embryo stage) (Meyer 2005). The maternal effect of the *mir-35* family suggests that these microRNAs function prior to the onset of sex determination, since the maternal lode of miRNA is established in the oocyte. Therefore, the *mir-35* family may not function simply to promote hermaphroditic fates in XX zygotes after sex determination has been established. Instead, we propose that expression of *mir-35* family miRNAs in the oocyte enables the *mir-35* family to function prior to the onset of sex determination to prevent the deleterious premature expression of sex-specific genes. The *mir-35* family is proposed to act upstream of its target genes *sup-26* and *nhl-2* in this process to prevent premature commitment to male programs. Accordingly, *sup-26* and *nhl-*2 are proposed to become relieved of *mir-35* family repression in XO embryos after the onset of sex determination, allowing them to promote the proper expression of male developmental programs (Figure 7G). This model of *mir-35* family function is consistent with the temporal expression pattern of the *mir-35* family, whose levels sharply decay in late-stage embryos (Stoeckius et al. 2009; Wu et al. 2010). Thus our model holds that a maternally-contributed miRNA family acts as a developmental timer, promoting naïveté in early embryos, and preventing the deleterious premature action of target RNAs involved in zygotic developmental decisions. This work therefore expands our understanding of miRNA functions and their contributions to developmental timing, and highlights the emergence of the *mir-35* family in particular as a model for understanding miRNA biology and mechanism prior to the OET.

The maternal effect of the *mir-35* family also implies that these microRNAs could function in both XX and XO embryos, since the maternal lode of miRNA is established in the oocyte, prior to the time that zygotic karyotype is established (at fertilization). Mutant *mir-35-41(nDf50)* XO animals display morphological defects in male-specific organs, which may be a result of the mis-timing of sex-specific gene expression (McJunkin and Ambros 2014). (i.e. While activation of male-specific gene expression *per se* would not be expected to perturb XO male development, the premature activation of these genes might disrupt the order of developmental events, and thus their fidelity.) Like XX embryos, XO embryos are affected by *mir-35* family mutant lethality (Figure S6). Whether the mis-timing of sex-specific gene expression or other developmental decisions is one of the molecular bases of this lethality is yet to be determined.

While the *mir-35* family’s maternal effect and maternal/embryonic expression indicate an early temporal window of *mir-35* family function, the spatial site of action of these miRNAs is less well understood. In this work, we show that SUP-26 and NHL-2 are both maternally inherited and localized to P granules in early embryos. We propose that aberrantly high levels of maternal or P lineage-localized SUP-26 and NHL-2 could be responsible for the cryptic Tra phenotypes observed in *mir-35* family mutants. This proposed site of action is consistent with our previous genetic studies which demonstrated that the *mir-35* family acts upstream of *sup-26* in the embryonic germline to ensure the temperature-robust fertility of adults (McJunkin and Ambros 2014). Aberrantly high expression of *her-1* in the P lineage of early embryos could be sufficient to drive a cryptic Tra phenotype since HER-1 acts non-cell-autonomously as a diffusible extracellular ligand (Hunter and Wood 1992).

SUP-26 was previously shown to modulate sex determination via translational control of *tra-2.* By generating a *sup-26(null)* allele, we have demonstrated that SUP-26 is not only a modulator of the pathway, but is in fact required for efficient male mating. In addition, the set of target genes bound by SUP-26 comprises a repertoire of many hundreds of mRNAs besides *tra-2*. In particular, SUP-26 binding of *dpy-30* could contribute to SUP-26’s regulation of *her-1* transcript via the DCC. Future studies should examine how SUP-26 exerts translational control on its many targets, and whether its function is always repressive or can include alternate modes of regulation.

Our work is the first indication that NHL-2 regulates sex determination. NHL-2 apparently acts upstream of *her-1* to promote male development through an unknown mechanism. NHL domains were recently shown to bind directly to RNA (Loedige et al. 2015); identifying all the RNA targets of NHL-2 will be critical in understanding its biological function in sex determination. In addition, the dramatic *mir-35-41(nDf50);her-1(gf)* synTra phenotype described here should enable forward genetic screens to identify downstream targets of NHL-2 and SUP-26 regulation.

The *mir-35-41(nDf50)* mutant gene expression pattern resembles that of the DCC component *sdc-2* that controls DC and sex determination. The majority of the similarities between these two mutants is attributable to expression changes related to perturbed sex determination, not direct effects of DCC(*lf*). Nonetheless, we observe that the most upstream regulator of sex determination, *her-1*, is derepressed in the *mir-35-41(nDf50)* mutant, implying that *mir-35-41* acts upstream of *her-1*, and possibly impacts the only known regulator of *her-1* transcription, the DCC itself. However, a primary hallmark of DCC(*lf*) would be XX-specific lethality due to impaired DC in somatic cells (Meyer 2005), yet the lethality of *mir-35* family mutants is not XX-specific (Figure S6). One possible explanation is that *sup-26* and *nhl-2* could be derepressed only in the P lineage of *mir-35* family mutant embryos, and not in the somatic lineages, where dosage compensation is critical for survival. Another possibility is that multiple pathways contribute to *mir-35* family mutant lethality, and that a form of karyotype-neutral lethality is epistatic to an underlying XX-specific lethality. It is also possible that *sup-26* and *nhl-2* may control the abundance of *her-1* in a DCC-independent manner.

The DCC is regulated by the competing zygotic expression of autosomal and X-linked genes, known as autosomal signal elements (ASEs) and X signal elements (XSEs), respectively. These signals converge to act on the master regulator XOL-1 (XO-Lethal 1), ensuring its low expression in XX embryos and high expression in XO embryos (Meyer 2005). XOL-1 prevents the expression of *sdc-2* - and thus DCC loading onto its target gene loci - in XOs. The exquisite gene dose dependence of *mir-35-41*, *sup-26*, and *nhl-2* in sex determination is consistent with these elements being involved in a counting mechanism like the competition between ASEs and XSEs. The *mir-35* family bears one characteristic of an XSE, in that it may promote DCC function. However, the *mir-35* family is not X-linked and has a strong maternal effect, both of which preclude its classification as an XSE. Intriguingly, in all other studied species of the *Caenorhabditis* clade, the *mir-35-41* cluster has undergone duplication events resulting in approximately twice as many copies of *mir-35* family miRNAs on X as on autosomes (*C. brenneri* 23 on X vs. 9 on autosomes, *C. remanei* 12 on X vs. 5 on autosomes, *C. briggsae* 16 on X vs. 9 on autosomes). Given the dosage-sensitive nature of this regulatory module, zygotic expression of the *mir-35* family members in other *Caenorhabditis* species may act as a counting mechanism to promote DCC loading or hermaphrodite sex determination specifically in XX embryos.

Like the *mir-35* family, small RNAs have been shown to play non-conserved roles in sex determination in *Bombyx mori*, *Drosophila melanogaster*, and *Zea mays* (Chuck et al. 2007; Kiuchi et al. 2014; Fagegaltier et al. 2014). The mechanism of action of small RNAs may suit them particularly well for involvement in sex determination pathways since sex determination is a rapidly-evolving process and the target specificity of small RNAs is also extremely fluid, likely due to the small genomic space that encodes the interaction interface – the 6-7 nucleotide seed (Shi et al. 2013). The *mir-35* family appears unique among these known mechanisms of sex determination by small RNAs because of its maternal effect on this zygotic developmental decision.

The *mir-35* family’s role in developmental timing is analogous to previously-characterized post-embryonically expressed miRNAs like *let-7* and *lin-4.* However, unlike *let-7* and *lin-4,* whose induction drives forward developmental transitions, the decay of the *mir-35* family members is likely to relieve repression of *sup-26* and *nhl-2*, allowing for the full onset of sex determination. For this reason, one might expect the timing of the decay of *mir-35* family members to be under precise developmental control. The sex determination pathway itself could potentially feed back on *mir-35-41*, possibly enforcing rapid decay in XO embryos, to permit the expression of male programs. How miRNAs are targeted for decay and turned over in the context of development is very poorly understood, and the newly-delineated *mir-35* network has the potential to illuminate such a process.

Whether repressed mRNAs can be re-expressed after decay of their cognate miRNA is another open question. In the embryo, miRNAs regulate the poly(A) tail length and translation of their targets (Wu et al. 2010; Bazzini et al. 2012; Subtelny et al. 2014). Since these mechanisms are theoretically reversible, transient miRNA-mediated repression followed by re-expression of the same mRNA molecule has the potential to occur in embryogenesis. This mode of regulation would be useful for storing maternally-contributed mRNAs in an inactive state for use later in development. Taken together, our work has immediate impact on our understanding of the biology of miRNA function and sex determination, and also lays the groundwork to address many important mechanistic questions regarding miRNA activity and the regulation of miRNAs.

## Materials and methods

### *C. elegans* culture and RNA preparation

*C. elegans* were cultured on HB101 *E. coli* by standard procedures at 20°C, unless otherwise specified. For 25°C microarray samples and *tra-2(ar221); xol-1(y9)* and *xol-1(y9)* samples, plates were upshifted to 25°C 24h prior to harvest. The samples for microarray analysis were grown on egg media. Embryos were isolated by hypochlorite treatment. All samples for RNA preparation were made by resuspending sample pellet in four volumes Trizol, snap freezing, and shaking for 15min at room temperature after thawing, prior to purification according to the Trizol manufacturer’s specifications (ThermoFisher). qRT-PCR was performed using Quantifast SYBR RT-PCR kit (QIAGEN). See supplemental experimental procedures for primer sequences, list of strains used, and details on vectors used for CRISPR.

### Microarrays and HITS-CLIP

Biotinylated antisense RNA was prepared using the Single-Round RNA Amplification and Biotin Labeling System (Enzo Life Sciences) and hybridized to Affymetrix 3’ IVT *C. elegans* microarrays. Expression data for *sdc-2(lf)* was acquired from NCBI GEO, Series GSE14649 (Jans et al. 2009). All data were analyzed using Genespring GX Software, excluding probes whose values were in the bottom quintile in every sample. CLIP was performed as in Zisoulis et al., with modifications as detailed in Supplemental Experimental Procedures. All microarray and HITS-CLIP data will be accessible at NCBI GEO, accession numbers XXXX.

### Immunofluorescence and immunoblot

Embryos were permeabilized by “freeze-cracking” (Duerr 2013) and fixed for 2min in methanol at −20°C then 30min at 25°C in 4% paraformaldehyde in PBS for staining with anti-FLAG antibody (M2 from Sigma Aldrich at 2μg/ml) or for 10min in methanol then 10min in acetone, both at −20°C, for staining with the anti-NHL-2 polyclonal (0.2μg/ml). Antibody staining was overnight at 4°C in 3% BSA with 0.1% Tween 20 and 0.1% Triton X-100 in PBS. P granule antibodies used were anti-PGL-1 (gift from C. Mello) diluted 1:500, and the K76 monoclonal diluted 1:100 (Strome and Wood 1983). Germlines were dissected and stained according to the Schedl lab protocols (http://genetics.wustl.edu/tslab/protocols/).

## Author Contributions

K.M. conducted the experiments. K.M. and V.A. designed experiments, interpreted results and wrote the paper.

## Acknowledgements

We thank the Walker lab for sharing their equipment, members of the Mello lab for helpful discussion and the anti-PGL-1 antibody, and C. Hammell for anti-NHL-2. The K76 P granule-specific antibody developed by S. Strome and W.B. Wood was obtained from the Developmental Studies Hybridoma Bank, created by the NICHD of the NIH and maintained at The University of Iowa, Department of Biology, Iowa City, IA 52242. Orkan Ilbay designed and cloned some of the vector variants used for genome editing. Thank you to J. Calarco for sharing modified CLIP protocols, to B. Meyer and R. Ketting for discussing unpublished data, and to A. Sitaram for critical reading of the manuscript. Some strains were provided by the CGC, which is funded by NIH Office of Research Infrastructure Programs (P40 OD010440). This work was funded by F32 GM097895, the Charles King Trust Postdoctoral Research Fellowship, and K99 GM113063-01 (to K.M.) and R01 GM34028 (to V.A.).

## References

Alvarez-Saavedra E, Horvitz HR. 2010. Many families of C. elegans microRNAs are not essential for development or viability. Curr Biol 20: 367–373.

Bazzini AA, Lee MT, Giraldez AJ. 2012. Ribosome profiling shows that miR-430 reduces translation before causing mRNA decay in zebrafish. Science 336: 233–7.

Chiang HR, Schoenfeld LW, Ruby JG, Auyeung VC, Spies N, Baek D, Johnston WK, Russ C, Luo S, Babiarz JE, et al. 2010. Mammalian microRNAs: experimental evaluation of novel and previously annotated genes. Genes {&} Dev 24: 992–1009.

Chuck G, Meeley R, Irish E, Sakai H, Hake S. 2007. The maize tasselseed4 microRNA controls sex determination and meristem cell fate by targeting Tasselseed6/indeterminate spikelet1. Nat Genet 39: 1517–1521.

Duerr JS. 2013. Antibody staining in C. elegans using “freeze-cracking”. J Vis Exp.

Fagegaltier D, König A, Gordon A, Lai EC, Gingeras TR, Hannon GJ, Shcherbata HR. 2014. A genome-wide survey of sexually dimorphic expression of Drosophila miRNAs identifies the steroid hormone-induced miRNA let-7 as a regulator of sexual identity. Genetics 198: 647–68.

Garcia DM, Baek D, Shin C, Bell GW, Grimson A, Bartel DP. 2011. Weak seed-pairing stability and high target-site abundance decrease the proficiency of lsy-6 and other microRNAs. Nat Struct Mol Biol 18: 1139–46.

Giraldez AJ. 2010. microRNAs, the cell’s Nepenthe: clearing the past during the maternal-to-zygotic transition and cellular reprogramming. Curr Opin Genet Dev 20: 369–75.

Greve TS, Judson RL, Blelloch R. 2013. microRNA control of mouse and human pluripotent stem cell behavior. Annu Rev Cell Dev Biol 29: 213–39.

Ha M, Kim VN. 2014. Regulation of microRNA biogenesis. Nat Rev Mol Cell Biol 15: 509–24.

Hammell CM, Lubin I, Boag PR, Blackwell TK, Ambros V. 2009. nhl-2 Modulates microRNA activity in Caenorhabditis elegans. Cell 136: 926–938.

Hammell M, Long D, Zhang L, Lee A, Carmack CS, Han M, Ding Y, Ambros V. 2008. mirWIP: microRNA target prediction based on microRNA-containing ribonucleoprotein-enriched transcripts. Nat Methods 5: 813–819.

Hargitai B, Kutnyánszky V, Blauwkamp TA, Steták A, Csankovszki G, Takács-Vellai K, Vellai T. 2009. xol-1, the master sex-switch gene in C. elegans, is a transcriptional target of the terminal sex-determining factor TRA-1. Development 136: 3881–3887.

Hunter CP, Wood WB. 1992. Evidence from mosaic analysis of the masculinizing gene her-1 for cell interactions in C. elegans sex determination. Nature 355: 551–5.

Hyenne V, Desrosiers M, Labbé J-C. 2008. C. elegans Brat homologs regulate PAR protein-dependent polarity and asymmetric cell division. Dev Biol 321: 368–78.

Jan CH, Friedman RC, Ruby JG, Bartel DP. 2011. Formation, regulation and evolution of Caenorhabditis elegans 3’UTRs. Nature 469: 97–101.

Jans J, Gladden JM, Ralston EJ, Pickle CS, Michel AH, Pferdehirt RR, Eisen MB, Meyer BJ. 2009. A condensin-like dosage compensation complex acts at a distance to control expression throughout the genome. Genes Dev 23: 602–18.

Kagias K, Pocock R. 2015. microRNA regulation of the embryonic hypoxic response in Caenorhabditis elegans. Sci Rep 5: 11284.

Ketting RF. 2011. microRNA Biogenesis and Function: An overview. Adv Exp Med Biol 700: 1–14.

Kiuchi T, Koga H, Kawamoto M, Shoji K, Sakai H, Arai Y, Ishihara G, Kawaoka S, Sugano S, Shimada T, et al. 2014. A single female-specific piRNA is the primary determiner of sex in the silkworm. Nature 509: 633–6.

Liu M, Liu P, Zhang L, Cai Q, Gao G, Zhang W, Zhu Z, Liu D, Fan Q. 2011. mir-35 is involved in intestine cell G1/S transition and germ cell proliferation in C. elegans. Cell Res 21: 1605–1618.

Loedige I, Jakob L, Treiber T, Ray D, Stotz M, Treiber N, Hennig J, Cook KB, Morris Q, Hughes TR, et al. 2015. The Crystal Structure of the NHL Domain in Complex with RNA Reveals the Molecular Basis of Drosophila Brain-Tumor-Mediated Gene Regulation. Cell Rep 13: 1206–20.

Mangone M, Manoharan AP, Thierry-Mieg D, Thierry-Mieg J, Han T, Mackowiak SD, Mis E, Zegar C, Gutwein MR, Khivansara V, et al. 2010. The landscape of C. elegans 3’UTRs. Science 329: 432–435.

Mangus DA, Evans MC, Jacobson A. 2003. Poly(A)-binding proteins: multifunctional scaffolds for the post-transcriptional control of gene expression. Genome Biol 4: 223.

Manser J, Wood WB, Perry MD. 2002. Extragenic suppressors of a dominant masculinizing her-1 mutation in C. elegans identify two new genes that affect sex determination in different ways. Genesis 34: 184–195.

Mapes J, Chen J-T, Yu J-S, Xue D. 2010. Somatic sex determination in Caenorhabditis elegans is modulated by SUP-26 repression of tra-2 translation. Proc Natl Acad Sci U S A 107: 18022–18027.

Massirer KB, Perez SG, Mondol V, Pasquinelli AE. 2012. The miR-35-41 family of microRNAs regulates RNAi sensitivity in Caenorhabditis elegans. PLoS Genet 8: e1002536.

McJunkin K, Ambros V. 2014. The embryonic mir-35 family of microRNAs promotes multiple aspects of fecundity in Caenorhabditis elegans. G3 (Bethesda) 4: 1747–1754.

Meyer BJ. 2005. X-Chromosome dosage compensation. WormBook 1–14.

Neumüller RA, Betschinger J, Fischer A, Bushati N, Poernbacher I, Mechtler K, Cohen SM, Knoblich JA. 2008. Mei-P26 regulates microRNAs and cell growth in the Drosophila ovarian stem cell lineage. Nature 454: 241–5.

Parchem RJ, Moore N, Fish JL, Parchem JG, Braga TT, Shenoy A, Oldham MC, Rubenstein JLR, Schneider RA, Blelloch R. 2015. miR-302 Is Required for Timing of Neural Differentiation, Neural Tube Closure, and Embryonic Viability. Cell Rep 12: 760–73.

Shi Z, Montgomery TA, Qi Y, Ruvkun G. 2013. High-throughput sequencing reveals extraordinary fluidity of miRNA, piRNA, and siRNA pathways in nematodes. Genome Res 23: 497–508.

Stoeckius M, Maaskola J, Colombo T, Rahn H-P, Friedländer MR, Li N, Chen W, Piano F, Rajewsky N. 2009. Large-scale sorting of C. elegans embryos reveals the dynamics of small RNA expression. Nat Methods 6: 745–751.

Strome S, Wood WB. 1983. Generation of asymmetry and segregation of germ-line granules in early C. elegans embryos. Cell 35: 15–25.

Subtelny AO, Eichhorn SW, Chen GR, Sive H, Bartel DP. 2014. Poly(A)-tail profiling reveals an embryonic switch in translational control. Nature 508: 66–71.

Ventura A, Young AG, Winslow MM, Lintault L, Meissner A, Erkeland SJ, Newman J, Bronson RT, Crowley D, Stone JR, et al. 2008. Targeted Deletion Reveals Essential and Overlapping Functions of the miR-17˜92 Family of miRNA Clusters. Cell 132: 875–886.

Wu E, Thivierge C, Flamand M, Mathonnet G, Vashisht AA, Wohlschlegel J, Fabian MR, Sonenberg N, Duchaine TF. 2010. Pervasive and cooperative deadenylation of 3’UTRs by embryonic microRNA families. Mol Cell 40: 558–570.

Zisoulis DG, Kai ZS, Chang RK, Pasquinelli AE. 2012. Autoregulation of microRNA biogenesis by let-7 and Argonaute. Nature 486: 541–544.

